# Feature-specific reaction times reveal a semanticisation of memories over time and with repeated remembering

**DOI:** 10.1101/2020.09.11.292813

**Authors:** Julia Lifanov, Juan Linde-Domingo, Maria Wimber

## Abstract

Memories are thought to undergo an episodic-to-semantic transformation in the course of their consolidation. We here tested if repeated recall induces a similar semanticization, and if the resulting qualitative changes in memories can be measured using simple feature-specific reaction time probes. Participants studied associations between verbs and object images, and then repeatedly recalled the objects when cued with the verb, immediately and after a two-day delay. Reaction times during immediate recall demonstrated that conceptual features were accessed faster than perceptual features. Consistent with a semanticization process, this perceptual-conceptual gap significantly increased across the delay. A significantly smaller perceptual-conceptual gap was found in the delayed recall data of a control group who repeatedly studied the verb-object pairings on the first day, instead of actively recalling them. Our findings suggest that wake recall and offline consolidation interact to transform memories over time, strengthening meaningful semantic information over perceptual detail.

## Introduction

One powerful way to protect memories against forgetting is to frequently recall them. Decades of research on the testing effect have shown such a protective effect, suggesting that repeated remembering stabilizes newly acquired information in memory (Abott, 1909; Gates, 1917; Roediger & Karpicke, 2006; Roediger & Butler, 2011; Dunlosky, Rawson, Marsh, Nathan, & Willingham, 2013). It is unknown, however, whether all aspects of a memory equally benefit from active recall. The aim of the present work was to investigate the qualitative changes in memories that occur with time and repeated remembering. We used feature-specific reaction time probes to measure such changes in lab-based visual memories. Specifically, we expected to observe a transformation along a detailed-episodic to gist-like-semantic gradient, based on several strands of research indicating that memories become “semanticised” in the process of their stabilisation.

Dominant theories of the testing effect make the central assumption that active recall engages conceptual-associative networks more so than other practice techniques like repeated study (Carpenter, 2011; Bjork, 1975; Kolers & Roediger, 1984). The elaborative retrieval account suggests that during recall, a conceptual relationship is established between initially separate episodic elements to unify them into a coherent memory (Carpenter, 2009). Similarly, the mediator effectiveness hypothesis (Pyc & Rawson, 2010) states that testing promotes long-term retention by evoking mediator representations, which are concepts that have meaningful overlap with a memory cue and target (Carpenter, 2011). Together, this work suggests that remembering co-activates semantically related concepts, more than restudy, and can thereby contribute to the long-term storage of newly acquired memories by linking them to already established, related concepts.

Other authors have made similar assumptions from a more neurobiologically and computationally motivated perspective (Antony, Ferreira, Norman & Wimber, 2017), drawing a parallel between the processes stabilizing memories via online recall, and the processes thought to consolidate memories via offline replay, including during sleep. In this online consolidation framework of the testing effect, active recall activates a memory’s associative index in the hippocampus, together with the neocortical nodes representing the various elements contained in the memory. As a result of this simultaneous activation, links between the active elements are strengthened (Hebb, 1949). Moreover, because recall tends to be somewhat imprecise, more so than re-encoding, activation spreads to associatively or conceptually related elements, providing an opportunity to integrate the new memory with related information. This presumed stabilization and integration is strongly reminiscent of the hippocampal-neocortical dialogue assumed to happen during sleep-dependent memory replay (Frankland & Bontempi, 2005), resulting in the integration of new memories into existing relational knowledge, and the strengthening of conceptual/schematic links between memories (Káli, & Dayan, 2004). Critically, many consolidation theories assume that this reorganization goes along with a “semanticisation” of memories, such that initially detail-rich episodic memories become more gist-like and lose detailed representations over time and with prolonged periods of consolidation (McClelland, McNaughton, & O’Reilly, 1995; Dudai, Karni, & Born, 2015; Winocur & Moscovitch, 2011). Based on these parallels between wake retrieval and offline consolidation, the present study tested whether repeated recall specifically induces a measurable “semanticisation” that goes beyond the effects that naturally occur over time.

In the human memory consolidation literature, much of the empirical evidence for semanticisation comes from neuroimaging studies showing a gradual shift in the engagement of hippocampus and neocortex during recent and remote recall, or tracking representational changes in memories over time (for a review, see Dudai et al., 2015; Tompary, Davachi, 2017). Behavioural studies have largely relied on scoring of autobiographical or other descriptive verbal memory reports for central gist versus peripheral details, and yielded robust evidence for a detail-to-gist gradient (e.g. Moscovitch, Cabeza, Winocur & Nadel, 2016; Sekeres, Bonasia, St-Laurent, Pishdadian, Winocur, Grady & Moscovitch, 2016). The present study used a different approach, asking if semanticisation via recall can be observed in reaction times (RTs) that specifically reflect the speed with which participants can access higher-level conceptual and lower-level perceptual features of visual object memories. This method was recently introduced by Linde-Domingo, Treder, Kerrén, & Wimber (2019). They showed that when participants are retrieving visual objects from memory, conceptual aspects (e.g., Does the recalled image represent an animate or inanimate object?) are accessed more rapidly than perceptual aspects (e.g., Does the recalled image represent a photo or a drawing?). In sharp contrast, RTs were consistently faster to perceptual than conceptual questions when the image was physically presented on the screen. This flip suggests that recalling a memory progresses in the opposite direction from visual perception, reactivating the core meaning first before back-propagating to sensory details. Such semantic prioritisation is plausible considering that the hippocampus is most directly and reciprocally connected with late sensory processing areas assumed to represent abstract concepts (Felleman & Van Essen, 1991; Suzuki & Amaral, 1994). Both online retrieval and offline replay of hippocampus-dependent memories can therefore be assumed to preferentially activate conceptual features of a memory, and this prioritisation may over time produce a semanticized memory compared with the one originally encoded. With this background in mind, and an adapted version of the described RT paradigm, we here investigate whether repeated retrieval enhances the semanticisation of memories over time compared to repeated study.

In our visual-associative learning experiment, two groups of participants were asked to learn novel verb-object pairings at the beginning of a first session (Fig. 1). They then immediately practiced those associations twice in each of three cycles, six times overall. Subjects in the retrieval group (n = 49) practiced by actively recalling the object image from memory when cued with the verb. Critically, in each of the three cycles they were asked to answer one conceptual and one perceptual question about the recalled object as fast as possible.

**Figure 1.**
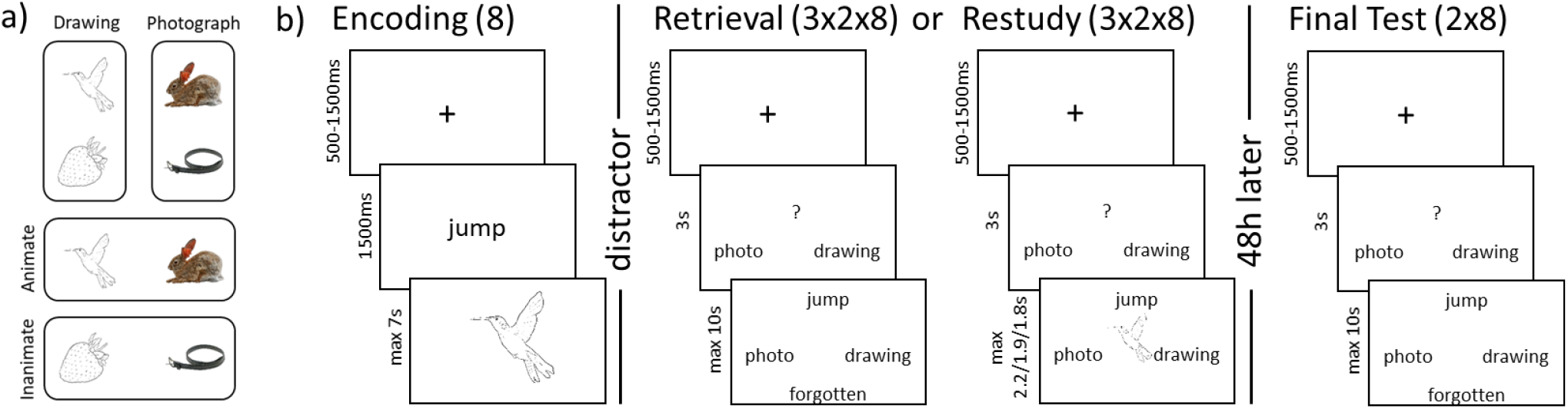
a) The design of the stimuli. The 64 pictures used in any given participant were orthogonally split into 32 drawings and 32 photographs, out of which 16 were animate and 16 inanimate objects, respectively. b) One prototypical task block of the paradigm within the repeated retrieval/restudy group. Both groups performed eight blocks, each starting with the encoding of eight novel verb-object associations. After a 20s distractor task, each of the eight associations was practiced twice in each of the three practice cycles, once with a conceptual, once with a perceptual question, and RTs were measured on each of the overall 6 practice trials. The maximum response time in each practice cycle of the restudy group was set to the average response time of the corresponding cycle in the retrieval group. After 48 hours, participants returned to complete a final test, where again each association was tested once with each of the two question types, with RTs being recorded. Finally, a written cued recall test was performed.

Subjects in the restudy group (n = 24) instead practiced by re-encoding the intact verb-object pairings, and were similarly asked to answer a conceptual and a perceptual question about the object in each cycle, but they did so while seeing the object on the screen. All participants returned to the lab 48h later for a delayed cued recall test, where each verb-object pairing was probed once more with a conceptual and once with a perceptual question.

We used reaction times (RTs) to the two types of questions as a measure of feature accessibility, probing the speed at which participants can access lower-level perceptual or higher-level conceptual features of the mentally reconstructed objects. We made two central predictions about how these RTs would change over time. First, we hypothesized that in the group who repeatedly recalled the associations immediately, the difference between perceptual and conceptual RTs will be significantly larger on the second testing day compared to the first day, reflecting time-dependent semanticisation. Second, we hypothesized that if retrieval plays a central role in this presumed semanticization, a control group who repeatedly studies the associations on the first day would show a significantly smaller perceptual-conceptual RT gap on the delayed memory test.

## Results

### Semanticisation over repeated retrieval

We first tested the retrieval group data for a time-dependent semanticisation, assuming that memory recall prioritises access to conceptual over perceptual features, and that this prioritisation increases over time with increasing semanticisation. We therefore tested whether the difference between conceptual and perceptual RTs is significantly larger on the second testing day compared to the end of the first day in the retrieval group, performing a 2 (recall cycle: end of day 1 vs day 2) by 2 (question type: conceptual vs perceptual) repeated measures analysis of variances (rmANOVA) on the RT data of the repeated retrieval group only (Fig. 2). This rmANOVA showed a main effect of recall cycle (F(1,48)=71.44, p<.01) indicating slower responses on day 2 than day 1, and a main effect of question type (F(1,48)=29.58, p<.01) with conceptual questions being consistently answered faster than perceptual questions. Critical to our first main hypothesis, the rmANOVA also revealed a significant interaction (F(1,48)=19.87, p<.01) between the two factors, indicating that the conceptual-over-perceptual RT advantage changed across days. A posthoc power analysis in G*Power revealed an effect size of d=.64 and a power of 0.99 for the interaction effect. Average RTs confirmed that the interaction was produced by an increasing perceptual-conceptual RT gap from day 1 (M_day1_ = .04, SD_day1_ = .19) to day 2 (M_day2_ = .29, SD_day2_ = .36), in line with the semanticization hypothesis.

**Figure 2.**
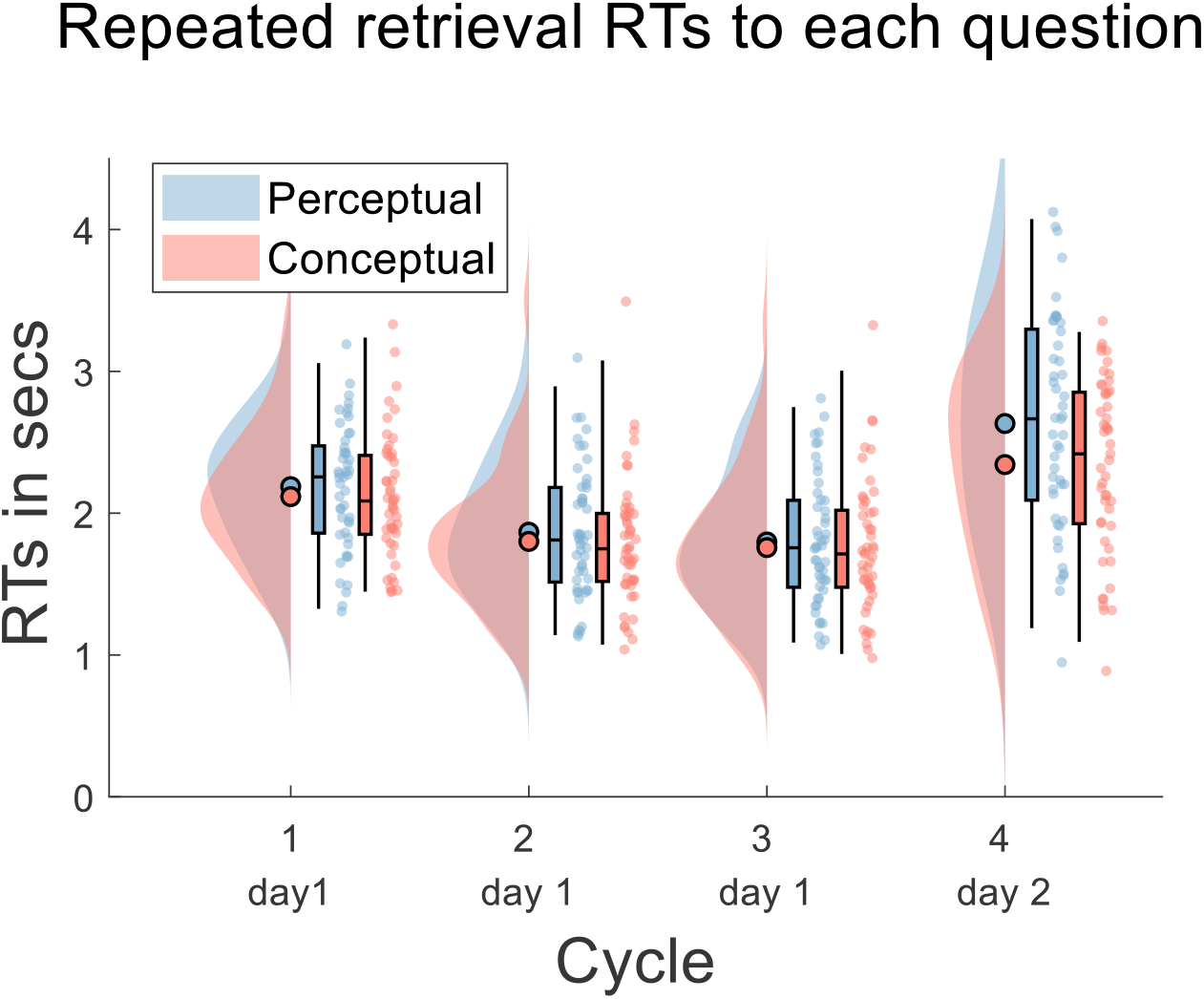
RTs in the repeated retrieval group for each cycle and question type. Filled circles represent the overall mean, boxplots represent median and quantiles, dots represent the means of individual subjects.

We additionally tested whether the perceptual-conceptual gap in the retrieval group changed across cycles on day 1, to explore whether these changes already take place over the first day, in line with a “fast consolidation” process (Antony et al, 2017). A 3 (cycle: 1, 2, 3) by 2 (question type: conceptual or perceptual) repeated measures ANOVA of the day 1 RTs (Fig. 2) revealed a significant main effect of cycle (F(2,96))=102.44, p<.01), with participants becoming faster over time, as well as a significant main effect of question type (F(1,48)=5.01, p=.03), with conceptual questions being answered overall faster than perceptual ones, generally replicating the results of Linde-Domingo et al. (2019). However, the cycle by question type interaction was not significant (F(2,96)=.42, p=.66), indicating that the perceptual-conceptual gap did not change significantly across practice cycles. The immediate recall data thus provide no evidence for a fast semanticisation.

### Semanticisation is facilitated more by repeated retrieval than restudy

To test our second hypothesis, that repeated retrieval leads to a stronger delayed perceptual-conceptual gap than repeated study, we investigated the RT gap on the second testing day in both groups. If semanticisation over time is enhanced by retrieval practice, this should be reflected in a larger RT gap in the retrieval group. A 2 (practice condition: retrieval vs restudy) by 2 (question type: conceptual vs perceptual) mixed ANOVA on the RTs of day 2 (Fig. 3) revealed no main effect of practice condition (F(1, 71) = 1.41; p = .24), and a main effect of question type (F(1, 71) = 16.92; p < .01) with shorter RTs for conceptual questions. As hypothesized, a significant interaction was found between question type and practice condition (F(1, 71) = 5.21; p = .03). Our posthoc power analysis on the interaction effect revealed an effect size of d=.27 and a power of 0.99. This interaction was due to an effect in the expected direction, with a higher perceptual–conceptual difference in the repeated retrieval group (M_retrieval_ = .29, SD_retrieval_ = .36) than in the restudy group (M_restudy_ = .08, SD_restudy_ = .37), in line with the interpretation that repeated retrieval leads to more pronounced semanticisation than repeated study.

**Figure 3.**
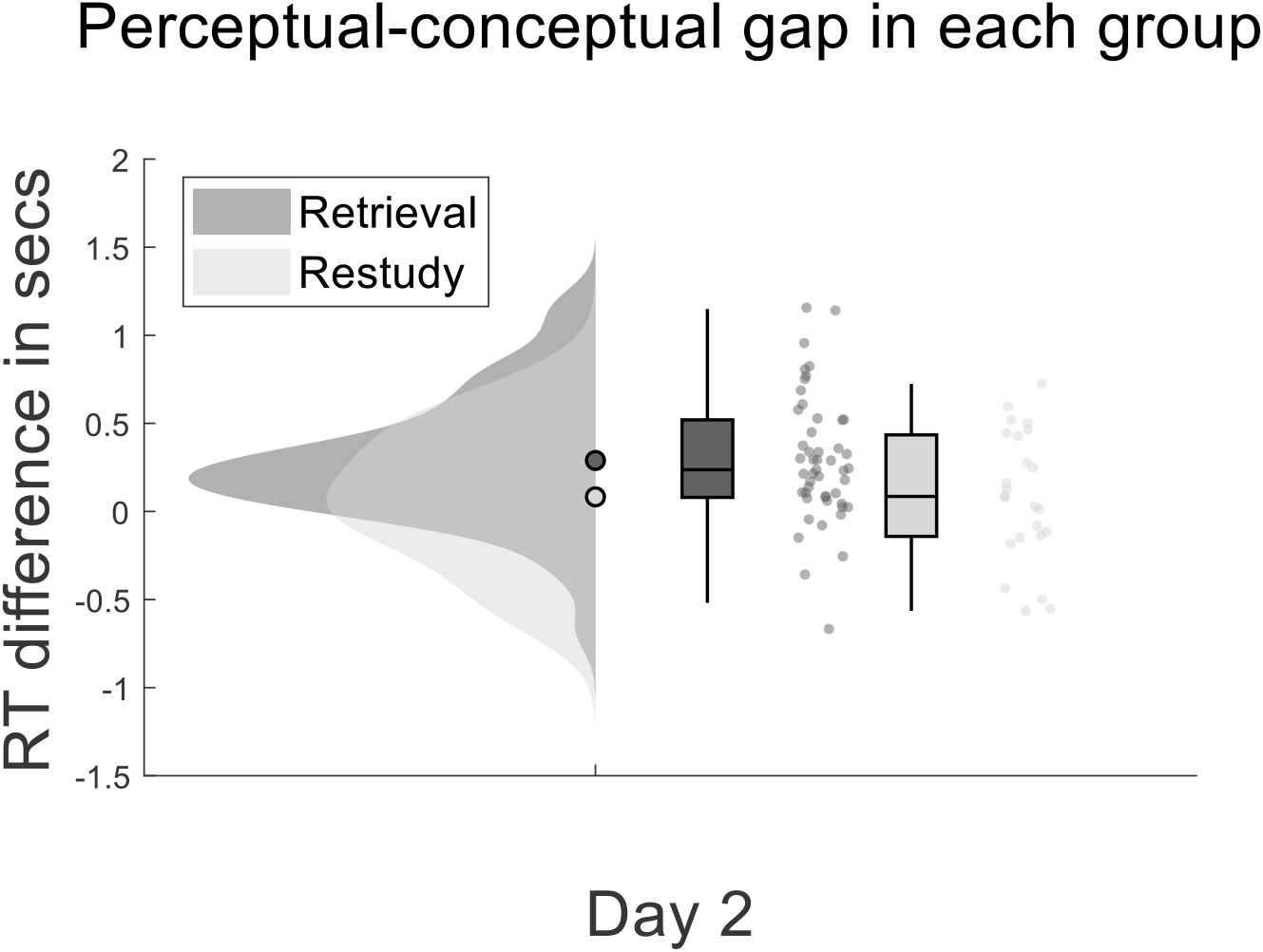
Perceptual-conceptual RT gap for both groups. Filled circles represent the overall mean, boxplots represent median and quantiles, dots represent the means of individual subjects.

### A replication of the reversed retrieval stream compared to restudy

Further, we analysed the data of the first day to test if we could replicate a reversal of the RT patterns between memory retrieval and visual exposure, conceptually replicating previous results (Linde-Domingo et al., 2019). Based on these findings, we expected faster RTs to conceptual than perceptual questions (i.e. a reverse stream) in the retrieval group (Fig. 2), and faster perceptual than conceptual RTs (i.e., a forward stream) in the restudy group. We therefore performed a mixed 2 (practice condition: retrieval vs restudy) by 2 (question type: conceptual vs perceptual) ANOVA on the data of day 1, averaging RTs across the 3 cycles. Apart from a main effect of task (F(1,71)=71.13, p<.01), and no main effect of question type (F(1,71)<.01, p=.98), this analysis revealed the expected, significant cross-over interaction (F(1,71)=9.24, p<.01) with faster responses for perceptual questions than conceptual ones in restudy (M_per_=1.13, SD_per_=0.21; M_con_=1.19, SD_con_=0.19) and vice versa in retrieval (M_per_=1.95, SD_per_=0.42; M_con_=1.89, SD_con_=0.44).

### Hierarchical relationship between remembered features

Two further analyses were conducted on accuracy data, rather than reaction times. First, we investigated a possible hierarchical dependency between perceptual and conceptual features as shown in recent work (Balaban, Assaf, Arad Meir & Luria, 2020), and how this relationship changed over time. All recall trials were sorted into four categories, depending on whether participants remembered both features, only perceptual features, only conceptual features, or none. In line with previous work (Balaban et al., 2020; Joensen, Gaskell & Horner, 2018), we expected that over time, the majority of items would be forgotten in a holistic manner, such that items that were fully remembered (“both features correct”) on day 1 would be fully forgotten (“none correct”) on day 2. For the present purpose, we were however particularly interested in the two response categories indicating partial remembering (i.e., “conceptual only” and “perceptual only” recall trials). Here, a hierarchical dependence in a reverse memory reconstruction stream predicts a particular pattern: higher-level conceptual information would need to be accessed before the lower-level perceptual information can be reached. As a result, participants should be relatively likely to remember the conceptual feature (“Was it animate or inanimate”) while forgetting the perceptual one (“Was it a photo or drawing”), but there should be very few trials where they remember the perceptual while forgetting the conceptual feature, except for random guesses. We thus expected to see a significant difference in the number of responses falling into these two categories already on the immediate day 1 recall. If semanticisation increases this hierarchical dependency, the gap in the proportion of conceptual-only and perceptual-only recalls should significantly increase across the 2-day delay.

We carried out a 2 (recall cycle: end of day 1 vs day 2) by 2 (features remembered: conceptual-only vs perceptual-only) rmANOVA to test this hypothesis. This analysis revealed a main effect of repetition (F(1,48)=55.52, p<.01), and a main effect of features remembered (F(1,48)=27.10, p<.01). Importantly, we also found the expected significant interaction (F(1,48)=8.21,p<.01), reflecting the observation that over time, the number of objects for which the conceptual but not the perceptual feature could be remembered increased significantly more than the number of objects for which the opposite pattern was true (Fig. 4).

**Figure 4.**
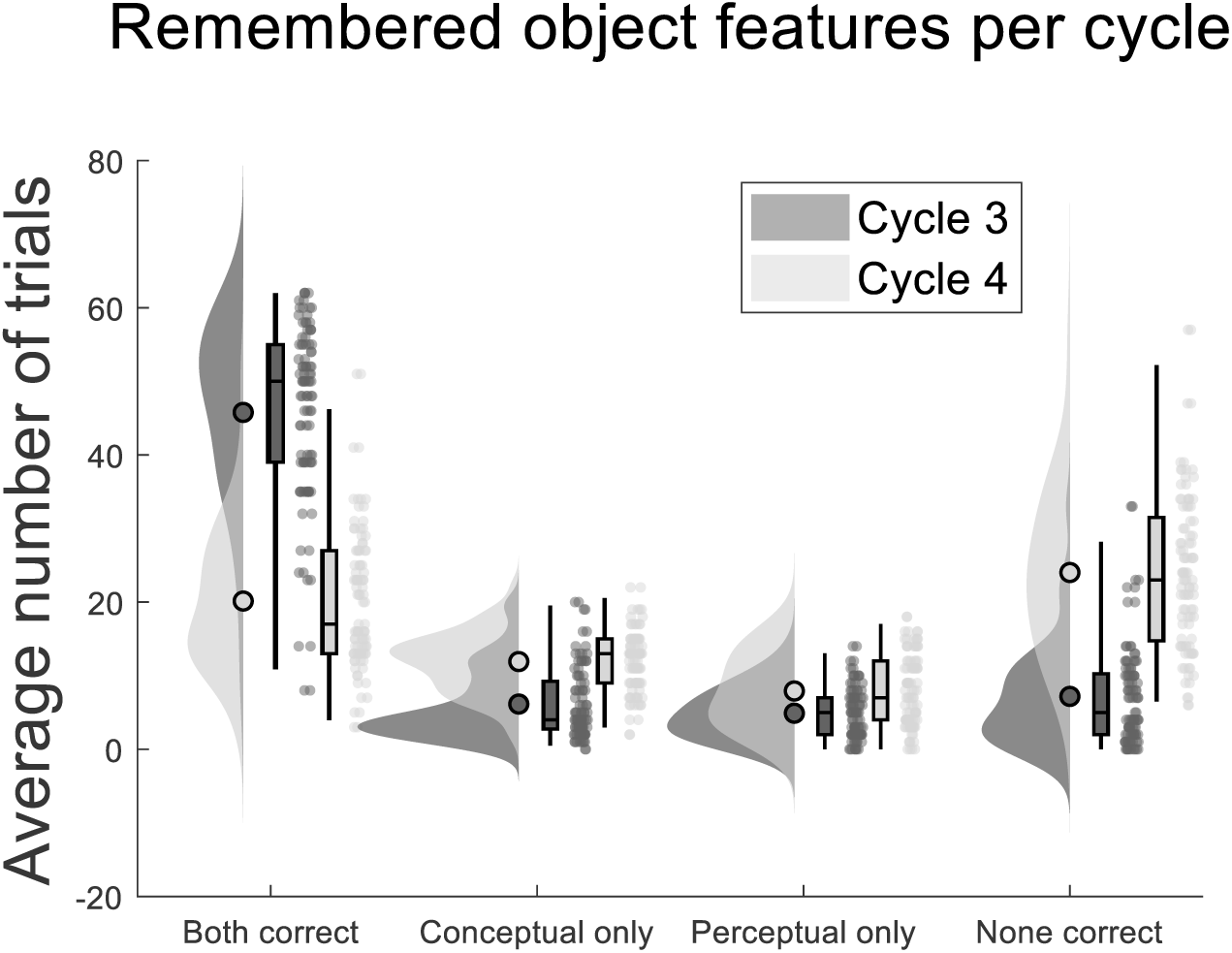
The average number of objects in each response-category for cycle 3 (end of day 1) and cycle 4 (day 2) of all subjects in the repeated retrieval group. Filled circles represent the overall mean, boxplots represent median and quantiles, dots represent the means of individual subjects.

Note that the data presented in the figure is not corrected for estimated random guesses (Balaban et al., 2020), as such a correction would have turned most proportions negative, and therefore seemed to be an overestimation of guesses in our dataset. However, since the guesses of a particular cycle are assumed to be distributed equally across response categories within that cycle, correcting will not change the outcomes of the statistical analysis.

### A replication of the testing effect

Finally, we also assessed the written cued recall responses on the second day to investigate if a general testing effect was found in our sample. To do so, we compared the accuracy in the written sheet responses between both experimental groups, using two independent sample t-tests. All written responses were categorized by two experimenters as “specific correct/incorrect” and “coarse correct/incorrect” responses. Here, specific correct includes retrieving the exact object label (e.g., parrot), whereas coarse correct responses also include correct descriptions of the object’s category (such as “colourful bird” for “parrot”). T-tests revealed that participants in the repeated retrieval group (M_coarse_=.30, SD_coarse_=.19; M_specific_=.25, SD_specific_=.18) recalled significantly more associations than restudy participants (M_coarse_=.20, SD_coarse_=.18; M_specific_=.16, SD_specific_=.17) using either scoring scheme, specific (T(71)=2.06, p=.04, CI= [.00, .17]) and coarse (T(71)=2.16, p=.03, CI= [.01, .19]). This finding confirms that our sample, not surprisingly, did show a testing effect in commonly used cued recall accuracies.

## Discussion

Do memories change every time we remember them? Cognitive (Carpenter, 2009; Pyc & Rawson, 2010) and neurobiologically motivated (Antony et al., 2017) theories assume that each active recall constitutes a distinct online consolidation event that systematically changes the nature of the memory, from an initially detail-rich episode to a “semanticised” version of the same event. Two questions were of central interest in the present study. First, we wanted to test if feature-specific probes can be used to reveal this presumed perceptual-to-conceptual transformation (semanticisation) of memories over an initial period of consolidation. Second, we were interested if repeated remembering specifically boosts this transformation compared with repeated study, preserving conceptual information relatively more over time.

To test our first hypothesis of a semanticisation over time, we measured how fast participants were able to recall perceptual and conceptual features of previously memorised objects on the first day, compared with how fast they accessed the same features on the second day. While conceptual information was consistently accessed faster, the perceptual-conceptual gap significantly increased over the two-day retention period, suggesting that access to conceptual memory features suffered less from the temporal delay than access to perceptual features. This finding is consistent with at least two possible interpretations. High-level semantic information may be prioritised for active consolidation, an ongoing discussion in the consolidation literature (Dudai et al, 2015; Schreiner & Rasch, 2018). Or semantic information might be forgotten at a slower rate than perceptual information, a possibility we return to further below. As also elaborated later on, hierarchical forgetting and prioritisation for active consolidation may in fact rely on the same underlying mechanism.

Recent studies do support an active and selective consolidation view. For example, structured, categorical information shows above-baseline enhancement from sleep, compared with detailed, stimulus-unique features of the memorized stimuli. It has thus been suggested that structured information is object to active consolidation (Schapiro, McDevitt, Chen, Norman, Mednick, Rogers, 2017). Because the hippocampus is most directly connected with late sensory areas coding abstract-semantic features of objects (Felleman & Van Essen, 1991; Suzuki & Amaral, 1994), the presumed hippocampal-neocortical dialogue during sleep might prioritise such conceptual features of the reactivated memories, relative to their perceptual features that are coded in brain areas further removed from the hippocampus. As a result, each replay event would strengthen semantic information more than perceptual, further exaggerating the gap that is already present on the immediate recall.

Alternatively, it is possible that the perceptual features of our visual objects were forgotten faster than their conceptual features. The nature of item-based forgetting is still under debate (Andermane, Joensen & Horner, 2020). Some recent work suggests that the forgetting of perceptual features, such as colour, is independent of, and occurs faster than, forgetting of higher-level conceptual features such as item state or exemplar (Brady, Konkle, Alvarez, Oliva, 2013; Utochkin & Brady, 2020). In contrast, other research has shown that object memories are forgotten in a more holistic manner, with an interesting hierarchically dependent forgetting of perceptual and conceptual features (Balaban et al., 2020). Inspired by this work, we investigated a possible hierarchical dependency of forgetting in our own accuracy patterns. We indeed saw evidence for asymmetrical recall, such that if participants only recalled one of the two features, they were more likely to remember the conceptual but not the perceptual feature than vice versa. This asymmetry significantly increased over the two-day delay, again indicating an increasing dependence of remembering on conceptual features. Together with the reaction time results, our findings therefore support a view of hierarchically dependent remembering and forgetting of single item features, with lower-level perceptual features having a higher likelihood of being forgotten independently of higher-level semantic features.

To distinguish the contribution of active retrieval to such time-dependent consolidation effects, we further tested whether retrieval on the first day enhances the preservation of conceptual features more than restudy, a more visual type of practice that does not involve the same degree of intrinsic memory reactivation. In line with our second main hypothesis, we found a larger perceptual-conceptual RT gap on the second day in the retrieval group, suggesting that the underlying semanticisation process is relatively stronger when the originally learned associations are immediately practiced by active cued recall. This finding has at least two important implications. First, sleep-dependent consolidation studies often carry out a memory test before and after sleep to obtain a difference score within subjects (Gais, Lucas & Born, 2006). Our results suggest that pre-sleep retrieval could be one driving factor for any representational changes that occur during the later overnight consolidation process (see also Cairney, Guttesen, El Marj & Staresina, 2018). Second, our finding has implications for theories of the testing effect, showing that active recall disproportionally strengthens conceptual aspects of a memory over perceptual ones. This finding resonates with the idea that each memory recall tests tend to co-activate semantically related information, in turn facilitating the integration of newly learned information into existing knowledge networks (Carpenter, 2009; Pyc and Rawson, 2010).

The time- and retrieval-induced memory changes found in the present study might be produced by neurocognitive processes that are shared between offline (replay-based) and online (retrieval-based) consolidation. It is well known that the hippocampus receives highly integrated, abstracted information from late sensory areas (Felleman & Van Essen, 1991; Suzuki & Amaral, 1994). Concept cells in the hippocampus are in fact assumed to form the building blocks of episodic memories (Quiroga, 2012). The various elements (e.g. objects, people) that constitute an episode are thus likely bound together on the level of meaningful semantic units. During retrieval (and offline replay), it is assumed that the linked elements belonging to the same episode are reactivated in a cascade that starts with pattern completion in the hippocampus, followed by a back-propagation into neocortex (Horner, Bisby, Bush, Lin & Burgess, 2015; Staresina & Wimber, 2019; Rolls, 2013). This back-propagation likely starts off with the information coded closest to the hippocampus, and then progresses backwards along the ventral visual hierarchy. If so, abstract-conceptual features of a memory should be reactivated first, faster than sensory-perceptual information. We previously provided evidence for this reverse mnemonic reconstruction from behaviour and decoding of brain activity patterns (Linde-Domingo et al., 2019), and we replicate this pattern in the present study. If abstract-semantic information is reactivated first during each offline replay or online retrieval event, it is more likely to benefit from these repeated reactivations, leading to a natural semanticization of memories over time.

Our findings support the idea that active testing, in terms of the neurobiological processes involved, mimics consolidation by relaying newly acquired information from hippocampal to neocortical structures (Antony et al., 2017; Ferreira, Charest & Wimber, 2019). However, the perceptual-conceptual gap in the retrieval group did not change with repeated remembering on day 1, and our results do thus not provide evidence for a “fast” consolidation process as suggested by Antony et al. (2017). They instead indicate that a major semanticisation process took place between the first and the second testing day, suggesting an interaction between active testing and time dependent consolidation. Retrieval thus seems to exert its effects most strongly when followed by a prolonged period of consolidation. This conclusion is in line with our present study, as well as previous behavioural (Butler & Roediger, 2007) and neural (Ferreira, et al., 2019) evidence. At this point, we can only speculate about the nature of this interaction. For example, an offline consolidation process might preferentially stabilize recently active pathways, thereby prioritising previously recalled memories (see Wilhelm, Diekelmann, Molzow, Ayoub, Mölle & Born, 2011, for related findings), resembling a tagging and capture mechanism (Redondo and Morris, 2011). Alternatively, online retrieval might act as a fast consolidation event with immediate effects, but these effects (e.g. neocortical integration) only become visible once the detailed hippocampal trace has decayed over time (Antony et al., 2017; Ferreira et al., 2019). Our findings do not support one or the other interpretation, and further investigation is needed to fully understand how wake remembering and offline replay interact.

The present findings suggest that reaction times, paired with questions that differentially probe access to specific mnemonic features, are sensitive to the presumed time- and recall-dependent transformation of relatively simple, visual-associative memories. Our feature-specific reaction time method thus complements other approaches that are commonly used to test for qualitative changes in memories. These include the scoring of autobiographical memories according to how much gist or detailed information subjects report (e.g. used in Moscovitch, Cabeza, Winocur & Nadel, 2016); recognition-based measures using familiarity as a proxy for gist, and recollection as a proxy for detail (Guran, Lehmann-Grube, & Bunzeck, 2020); and more recently, measures of access and precision (Berens, Richards & Horner, 2020; Cooper & Ritchey, 2020). Reaction times are rarely used in memory studies. Object recognition work, however, shows that the speed with which participants can categorize objects (e.g., animate/inanimate) is well aligned with the time points when the same categories can be decoded from brain activity (Carlson, Ritchie, Kriegeskorte, Durvasula & Ma, 2013; Ritchie, Tovar & Carlson, 2015), and a recent study used the same approach to track the back-propagation of information during memory recall, suggesting that conceptual features are reached earlier during the reverse reconstruction than perceptual features (Linde-Domingo et al., 2019). The present results indicate that RTs can directly tap into the qualitative changes that occur over the course of memory consolidation.

In summary, using feature-specific probes, we provide evidence for the semanticisation of memories over time and specifically with repeated remembering. Our main results are consistent with a framework where the natural prioritisation of conceptual information during repeated retrieval (Linde-Domingo et al., 2019) has a lasting effect on what is being retained over time. We reconcile cognitive theories of the testing effect with neurobiologically motivated theories of memory retrieval, which posit that functional anatomy during retrieval dictates faster access to later and more conceptual stages of visual processing. Finally, our feature-specific RT probes provide a new alternative to assess the qualitative changes of mnemonic representations over time, and might thus be useful for future consolidation studies using lab-based rather than autobiographical memories.

## Methods

### Participants and a priori power calculations

Previously published work has found an effect size of d=.55 for the perceptual-conceptual gap in RTs during retrieval (Linde-Domingo et al., 2019). We expected an effect size at least as large on day 2 in the repeated retrieval group. A power analysis in G*Power (Faul, Erdfelder, Buchner, & Lang, 2009) with d=.55, α= .05 and a power of 0.9 suggested that a sample size of at least 30 was required to detect an existing effect in the retrieval group. The effect of most interest in the retrieval group was a significant interaction between testing day and question type, specifically such that the gap between conceptual and perceptual RTs would significantly increase from day 1 to day 2. The power for this interaction contrast could not be estimated a priori from the work of Linde-Domingo et al. (2019). To have sufficient power to detect an increase in the perceptual-conceptual gap, we decided to double their sample size, aiming for 48 subjects in the retrieval group (see results section for corresponding posthoc power analyses).

The second comparison of interest in this study was a contrast between the perceptual-conceptual gap on day 2 (i.e., delayed test) in the retrieval and the restudy groups. Again, since the effect size could not be estimated directly from previous work, we aimed for n=24 participants in the restudy group in order to reach a sample size of n=72 overall for the critical comparison of the retrieval and the restudy group. Posthoc power analyses can be found in the results section.

Fifty-seven healthy volunteers from the local student population in Birmingham participated in the retrieval condition (45 female and 12 male, mean age [M] = 19.95, standard deviation [SD] =.79), of which eight were excluded due to absence on the second testing day or missing data. Another 26 volunteers participated in the restudy group (21 female and 5 male, M =18.92, SD =.89), of which two were excluded due to absence on the second testing day. Our final sample thus consisted of 49 participants in the retrieval group and another 24 participants in the restudy group. All participants were informed about the experimental procedure, underwent a screening questionnaire (including sleep and consumption behaviour 24h before the experiment) and gave their written informed consent. The research was approved by the STEM ethics committee of the University of Birmingham.

### Material

The paradigm was an adapted version of the visual verb-object association task designed by Linde-Domingo et al. (2019). Our stimulus materials consisted of 64 action verbs and 128 pictures of everyday objects, all presented on white backgrounds (see Fig. 1.a; for more detailed information about the source of the verbs and pictures, see Linde-Domingo et al. (2019)). Importantly, objects were categorized into two conceptual classes, i.e. animate vs inanimate objects; and two perceptual classes, i.e. black line drawings vs coloured photographs. We pseudo-randomly drew 64 images per participant according to a fully balanced scheme, such that each of the two-by-two categories included the same number of pictures (16 animate-photographs, 16 animate -drawings, 16 inanimate-photographs, 16 inanimate-drawings). Action verbs were randomly assigned to images in each participant, and were presented together with pictures centrally overlaid on a white background. The stimulus presentation and timing and accuracy information collection was controlled by scripts written in Matlab 2017a (www.mathworks.com) and the Psychophysics Toolbox extension (Brainard, 1997; Pelli, 1997; Kleiner, Brainard, Pelli, Ingling, Murray & Broussard, 2007).

For the analysis we used customized Matlab code (https://www.mathworks.com/matlabcentral/fileexchange/64980-simple-rm-mixed-anova-for-any-design; https://www.mathworks.com/matlabcentral/fileexchange/6874-two-way-repeated-measures-anova). Figures were created using the raincloud plots tool (Allen, M., Poggiali, D., Whitaker, K., Marshall, T. R., & Kievit, R. A, 2018; Allen et al., 2019) and colorbrewer schemes for Matlab (https://www.mathworks.com/matlabcentral/fileexchange/34087-cbrewer-colorbrewer-schemes-for-matlab).

### Procedure overview

In both experimental groups, participants were informed about the experimental procedure, asked to sign an informed consent form, and to perform a training run. After completion of this training, participants continued to the experimental task (Fig 1.b). On day 1, participants performed eight task blocks, each including an encoding block with eight trials, a 20s distractor task and three practice cycles, each including two times eight practice trials. Returning after 48h, participants finished the experiment with a final test consisting of a single retrieval cycle (see below for details). Before leaving, participants completed a written cued recall test. It took participants about 70 min to perform the task on day 1, and about 20 min on day 2.

### Encoding

In each encoding block (Fig. 1.b), participants were instructed to study 8 novel verb-object pairings. A fixation cross was presented to the participants for a jittered time period between 500 and 1500ms. An action verb was then presented for 1500ms before an object was shown for a maximum time period of 7s. To facilitate learning, participants were instructed to form a vivid visual mental image using the verb-object pairing. Once they had formed a strong mental image, participants were asked to press the up-arrow key, which moved the presentation on to the next trial. In the repeated retrieval group, it took participants 4.65s on average, and in the restudy group it took them 4.34s to proceed to the next trial (SD_retrieval_=1.77; SD_restudy_=1.65).

### Distractor

After each encoding block, participants performed a self-paced distractor task for 20s, indicating as fast as possible whether each of the consecutively presented numbers on the screen was odd or even, using a left/right key press. Feedback on the percentage of correct responses was provided at the end of each distractor phase.

### Practice

#### Repeated retrieval group

The retrieval trials started with the presentation of a fixation cross, jittered between 500 and 1500ms, and followed by the conceptual (animate/inanimate) or perceptual (photo/drawing) question that was displayed for 3s, enabling participants to mentally prepare to recall the respective feature of the object that was relevant on a given trial. The verb was then displayed above the response alternatives (e.g., animate/inanimate), and participants had to retrieve the associated object and answer the question as fast as possible. Verb and question were displayed for a maximum period of 10s or until the participant selected a response to the question. The questions were answered with left, downward and right-arrow keys.

#### Restudy group

In the restudy group, the paradigm was kept as similar to the repeated retrieval group as possible, including an attempt to equate average exposure times during practice (for which reason the restudy group data was collected after the retrieval group). The restudy trial was initiated with a fixation cross with the same jitter (500-1500ms) as in the retrieval group, and followed by the conceptual or perceptual question that was displayed for 3s. The verb cue and object then appeared together above the question. Again, participants were asked to use the 3s period to prepare mentally to answer the question. When the object appeared, participants were instructed to first answer the question about the object they saw on the screen as fast as possible, and then use the remaining time to restudy the verb-object pair. In order to equate exposure times between the two groups, we set the trial duration of each of the three restudy cycles to the average response time of each of the three individual retrieval cycles from the previously collected retrieval group (cycle 1: 2.2s, cycle 2: 1.9s, cycle 3: 1.8s).

### Retrieval and restudy blocks setup

Participants of both groups completed three consecutive practice cycles, in each of which they practiced all eight verb-object associations they had learned in the previous encoding block twice, once answering a conceptual and once answering a perceptual question. This sums up to six practice trials per learned association, three with each question type. The order of the conceptual and perceptual questions within cycles was counterbalanced as follows: In each of the three cycles, one half of the stimuli was first probed with a conceptual question and the other half with a perceptual question first. Additionally, we controlled that each of the eight question-order possibilities occurred equally often for each object type (i.e., animate-photo, animate-drawing, inanimate-photo, inanimate-drawing). The percentage of correct trials was provided after the third practice cycle.

### Final Test

After 48 hours, participants were asked to complete a final test, in which they performed one cued recall block with the same procedural set-up as on day 1 in the retrieval group. Participants were presented with a conceptual/perceptual probe, and asked to answer this question as fast as possible when cued with a verb. Each object was recalled once with each question type. Here, half of the stimuli was first probed with a conceptual question and the other half with a perceptual question, randomized independently with respect to the first testing day. Finally, participants were given a paper sheet, displaying all 64 action verbs, next to which they were asked to write down a verbal description of the associated object.

### Data Preparation

During data preparation, RTs of correct trials were averaged and the standard deviation was calculated for both conceptual and perceptual questions, separately for the retrieval and the restudy group, and separately for the trials of each individual practice cycle per subject. Trials faster than 200ms, or exceeding the average RT of a given cycle by more than three times the standard deviation, were excluded in further RT analyses (Linde-Domingo et al., 2019). In the repeated retrieval group, 98.16 % of the data remained after trimming the RTs of correct responses, whereas in the restudy group, 99.60% remained.

To prepare the accuracy data, trials with responses faster than 200ms and objects with a no-response for either of both questions on one cycle were excluded in the related cycle. After this accuracy trimming, 99.39% of the repeated retrieval data and 93.26% of the restudy data remained. The RT data prepared for our main hypotheses met the normality assumptions.

### Data and code availability statement

The data and code that support the findings of this study are available under https://osf.io/wp4fu/?view_only=273c9f31c9464135a19b471e34b2330a.

